# Predicting perturbation patterns from the topology of biological networks

**DOI:** 10.1101/349324

**Authors:** Marc Santolini, Albert-László Barabási

## Abstract

High-throughput technologies, offering unprecedented wealth of quantitative data underlying the makeup of living systems, are changing biology. Notably, the systematic mapping of the relationships between biochemical entities has fueled the rapid development of network biology, offering a suitable framework to describe disease phenotypes and predict potential drug targets. Yet, our ability to develop accurate dynamical models remains limited, due in part to the limited knowledge of the kinetic parameters underlying these interactions. Here, we explore the degree to which we can make reasonably accurate predictions in the absence of the kinetic parameters. We find that simple dynamically agnostic models are sufficient to recover the strength and sign of the biochemical perturbation patterns observed in 87 biological models for which the underlying kinetics is known. Surprisingly, a simple distance-based model achieves 65% accuracy. We show that this predictive power is robust to topological and kinetic parameters perturbations, and we identify key network properties that can increase up to 80% the recovery rate of the true perturbation patterns. We validate our approach using experimental data on the chemotactic pathway in bacteria, finding that a network model of perturbation spreading predicts with ~80% accuracy the directionality of gene expression and phenotype changes in knock-out and overproduction experiments. These findings show that the steady advances in mapping out the topology of biochemical interaction networks opens avenues for accurate perturbation spread modeling, with direct implications for medicine and drug development.

**Significance statement:** The development of high-throughput technologies has allowed to map a significant proportion of interactions between biochemical entities in the cell. However, it is unclear how much information is lost given the lack of measurements on the kinetic parameters governing the dynamics of these interactions. Using biochemical networks with experimentally measured kinetic parameters, we show that a knowledge of the network topology offers 65% to 80% accuracy in predicting the impact of perturbation patterns. In other words, we can use the increasingly accurate topological models to approximate perturbation patterns, bypassing expensive kinetic constant measurement. These results could open new avenues in modeling drug action, and in identifying drug targets relying on the human interactome only.

## Introduction

In the past decade, we witnessed great progress towards the systematic and comprehensive mapping of the physical interactions between the biochemical entities that make up the cells, that together represent the human interactome (1–3). These advances have fueled the rapid development of network biology as a suitable framework to describe and understand cellular processes and how their collective perturbations affect disease states (1, 4–8). The underlying data, fueling these advances, include but are not restricted to protein-protein interactions, gene regulation, metabolic reactions or kinase-substrate interactions. Through the aggregation of systematic and literature derived interactions from multiple resources, the human interactome covers today 170,000+ physical interactions between ~14,000 biochemical entities (7). Complementing this wealth of the interaction data, a massive amount of data is routinely generated at the microRNA (miRNA), messenger RNA (mRNA) and protein level through large scale measurements of their abundance in various cell types, organisms and conditions, populating databases such as Gene Expression Omnibus (9).

From the knowledge of the interactome, the goal of network biology is to quantify and predict the spread of perturbations across the subcellular network. Such perturbation patterns are of crucial importance for network medicine, helping us understand the differential expression patterns observed in disease states. Moreover, being able to prioritize the effect of biological perturbations *in silico* is key given the cost, time and difficulty to obtain such data through perturbation experiments, especially for human subjects. Yet, while the coverage of the interactome is increasing steadily, network biology continues to lack a general quantitative dynamical framework for such predictive modeling. This is in part due to the rarity of large scale measurements of the kinetic parameters, necessary to populate the kinetic models of all pathways (10, 11). Moreover, the degree of reproducibility of the measured kinetic parameters varies wildly between *in vitro* and *in vivo* experiments (12). Other approaches pertaining to the global fitting of kinetic parameters (13, 14) by optimizing the model agreement to available data often yield large parameter uncertainties (15, 16). To bypass the need of a full knowledge of kinetic parameters, other studies have investigated the interplay between generic dynamical models and topological structure in the context of biological networks (17, 18). These studies have focused on retrieving global perturbation statistical properties from microscopic models (17), or retrieving the most probable underlying dynamical model from perturbation statistics (18). However, such universal insights are of limited predicting power when confronted with small-size biological models with heterogeneous dynamics. On the other hand, topological models have been proposed to study perturbation spread in biological networks, such as Boolean networks (19) or Normalized Hill models (20). However, such studies are usually limited to a few well described, small-size networks, not offering a comprehensive picture of the accuracy of topological models when applied to a large diversity of real-world biological networks.

While we continue to lack large scale measurements of kinetic parameters, the literature has flourished with detailed biochemical models of smaller scale. This is indicated by the growing body of databases dedicated to the storage of biological models (21). For example, the repository of computational models of biological processes BioModels has seen a steady growth of its content over the last decade and currently hosts over 1,200 models derived directly from the literature (22). These models contain detailed information on the pertinent biological components, their interactions and the differential equations describing their dynamics.

The fine-grained level of detail these biochemical models offer opens an avenue to explore what level of description is necessary (and sufficient) to reproduce the perturbation patterns characterizing biological networks. In particular, previous work has shown that some perturbation patterns are robust to significant changes in kinetic parameters (23, 24). Indeed, only a small subset of these model parameters affect the overall dynamics, a property known as “sloppiness” (25). Accordingly, a simplified dynamical model containing only a few parameters has been found to accurately predict perturbation patterns obtained with a full biochemical model in the case of the beta-adrenergic pathway (20). While these studies focus on perturbations around a given steady-state, their results have been extended to the full dynamical landscape of biological networks (26). Using random kinetic models, Huang et al. (26) have shown in the cases of toggle-switch-like motifs and a 22-node biological network that the stable states converge to experimentally observed gene state clusters even when the parameters are strongly perturbed, suggesting that the dynamics is determined mainly by the circuit topology, not by detailed kinetic parameters.

Here we develop DYNamics-Agnostic Network MOdels (DYNAMO), an ensemble of perturbation propagation models that rely on the network topology alone, and investigate the extent to which the relative magnitude of biological perturbations can be retrieved when we lack knowledge of the kinetic parameters and the details characterizing the dynamics of the underlying biochemical process. Using an ‘onion-peeling’ strategy across a variety of detailed biochemical models, we systematically quantify the loss of accuracy of the predicted perturbation patterns when we successively remove information on the specifics of the dynamics.

We show that an accurate knowledge of the network topology captures on average 65% of the influence patterns of the full biochemical model. We then identify global network features that guarantee higher accuracies, reaching up to 80% predictive power. The underlying modeling framework, and its level of accuracy are finally validated using perturbation experiments in the chemotaxis network in bacteria.

## Results

### 1. Modeling influence patterns in biological networks

When the concentration of a biological species is perturbed, the perturbation can spread along physical interactions and reactions, reaching other parts of the interactome (Figure 1a). A purely topological approach predicts a uniform spread across the network: first neighbors are affected the most, followed by second neighbors etc. In reality, each interaction is governed by a specific dynamical equation with an associated set of parameters, allowing for a precise computation of influence propagation. We must consider the full dynamics to determine the precise direction and the rate at which a perturbation spreads within the network.

**Figure 1.**
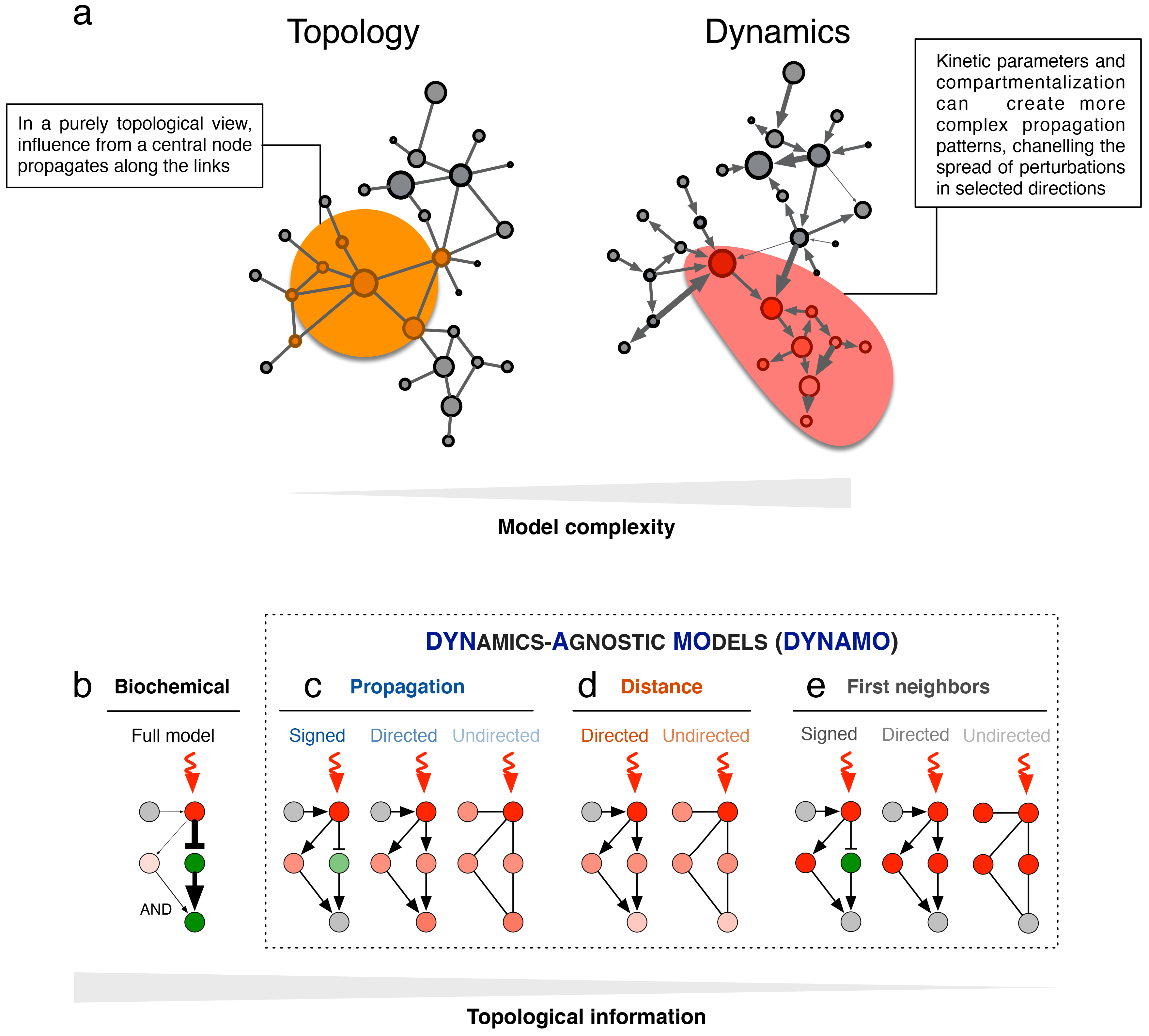
Influence patterns in biological networks. **a.** The multiple interactions between cellular components form the subcellular network or interactome. Mapping out the connections between these entities is the necessary first step to understanding how perturbations propagate in the network. On top of the network structure, the dynamics is obtained when adding the knowledge of link direction, sign, and kinetic parameters. The resulting complete biochemical model offers the best predictive model of influence propagation. b-e Schematic representations depicting the propagation of a perturbation in a biological network according to different DYNAMO models of decreasing complexity. Detailed description of the models can be found in the Methods section.

Here we investigate the degree to which these dynamical patterns can be retrieved from simple topological models. To that aim, we explore several models of influence propagation with increasing complexity, both in terms of accuracy of representation of the underlying network topology and in terms of dynamical model used (see Figure 1b-e and Methods). We describe network topology in terms of four layers of increasing complexity: (1) undirected network, (2) directed network, (3) directed and signed (activating/inhibiting) network, and (4) directed, signed and weighted network. To illustrate these layers, consider two interacting species A and B, where an increase in A causes B to increase while a change in B does not affect A. The four layers of complexity successively describe the knowledge that (1) A and B interact (the existence of a link), (2) A causes a change in B’s concentration (direction of influence), (3) A causes a positive change in B’s concentration (sign) and (4) A causes a positive change in B’s concentration of a certain strength (magnitude determined by the link weight parameters, like kinetic constant). All this information can be extracted from the Jacobian matrix of the system (see Methods), which quantifies the degree to which a change in A’s concentration causes a change in B’s concentration, and the direction of the change (positive for an increase or negative for a decrease). In this work, the Jacobian matrix is constructed from the underlying systems of dynamical equations characterizing a biological model. The signed Jacobian matrix is then used to reconstruct the underlying weighted topology of the biological models, allowing us to capture topologies of type (1) to (3). The topology of type (4), built from the full Jacobian matrix, contains the kinetic parameters information and hence corresponds to the full biochemical model. Therefore, from the Jacobian matrix, we extract both the topology and perturbation dynamics of the studied models.

Given a network topology (wiring diagram), we wish to predict how the perturbation of a given species propagates over the network and the degree to which it affects all other species. Such perturbation patterns – that we call *influence patterns* – are usually represented by a sensitivity matrix - also called the linear response matrix or correlation matrix in the literature (27)–, describing the change in the steady state value *x*_*i*_ of a node *i* when the steady-state value *x*_*j*_ of another node *j* is varied (27, 28): 
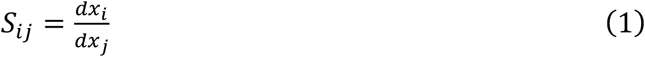

If the dynamical equations are known, the sensitivity matrix can be analytically derived using a perturbative framework (17, 27) (see Methods). In the following we refer to this exact sensitivity matrix as the “(full) biochemical model”, which is the underlying model from which it is computed (Figure 1b).

We explore three models of decreasing complexity to compute the sensitivity matrix using topological information only (Figure 1c-e). We refer to them as Dynamics-Agnostic Models or DYNAMO:

i. We start with a “propagation model” (Figure 1c) proposed in the context of disease gene prioritization, where “influence” spreads from a set of known seed genes to highlight putative disease genes (29). In our case, perturbed species are seed genes and we want to prioritize the perturbation level of the other species. In this model, the predicted perturbation of a node is proportional to the degree-weighted sum of the perturbations of its neighbors, with a constant input term added for the “source” node being perturbed. This propagation model has been shown to outperform a Random Walk algorithm in prioritizing disease genes across 1,369 diseases (29).
ii. The “distance model” (Figure 1d) assumes that the strength of a perturbation is inversely proportional to the network distance between a species and the source of perturbation, that is to the number of interactors it takes for one species to affect the expression of another. Such a model is of great interest since network distance in the interactome is a remarkable predictor of similarity between diseases (7) and drug-disease association (30).
iii. The minimal “first neighbor” model (Figure 1e) assumes that the perturbation reaches only the direct neighbors of a perturbed node. Such direct neighbor influence, also called the “local impact hypothesis” (31), has proven fruitful for disease gene prediction (32), and is at the core of the Minimum Dominating Set (MDS) controllability approach, where a minimal group of nodes is identified such that all other nodes in the network have a direct interaction with an MDS node and can therefore be suitably “controlled” or influenced. Applied to the protein-protein interactome, the resulting MDS proteins were shown to be more essential and disease-related that non-MDS proteins (33). As perturbations first impact the first neighbors of the perturbed nodes, the first neighbor model is the most minimal approximation we explore.

In the following we apply these models to a diverse set of well-characterized biochemical models.

### 2. Topological models accurately predict influence patterns in biological models

To test the validity of our findings on a large and diverse set of biological networks, we start from the BioModels database, a repository of curated biological, dynamical models (see Methods). Biological models from this database are deposited in a standard format allowing us to extract the underlying set of differential equations describing their dynamics using libSBML (34) (Figure 2a). From the dynamical equations, we derive the influence networks by linking a species *i* to a species *j* if a permanent change in the concentration of *i* directly affects the steady state concentration of species *j*. To do so, we first compute the Jacobian matrix of the system. Writing the dynamical equations as 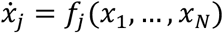, where *N* describes the number of species in the model and *x*_*i*_ is the concentration of species *i*, the Jacobian is 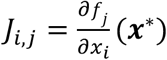, where ***x***\* is the steady-state of the system (Figure 2b). The adjacency matrix *A* of the network is computed as *A* = *sign*(*J*^*T*^(*x*\*)), where *J*^*T*^denotes the transpose of the Jacobian of the system and the sign function applies element-wise (see Methods and Figure 2c). In this framework, links with negative weights correspond to inhibitory interactions while positive weights denote activating interactions. This approach is reminiscent of inference networks (35), with link directions systematically reversed.

**Figure 2.**
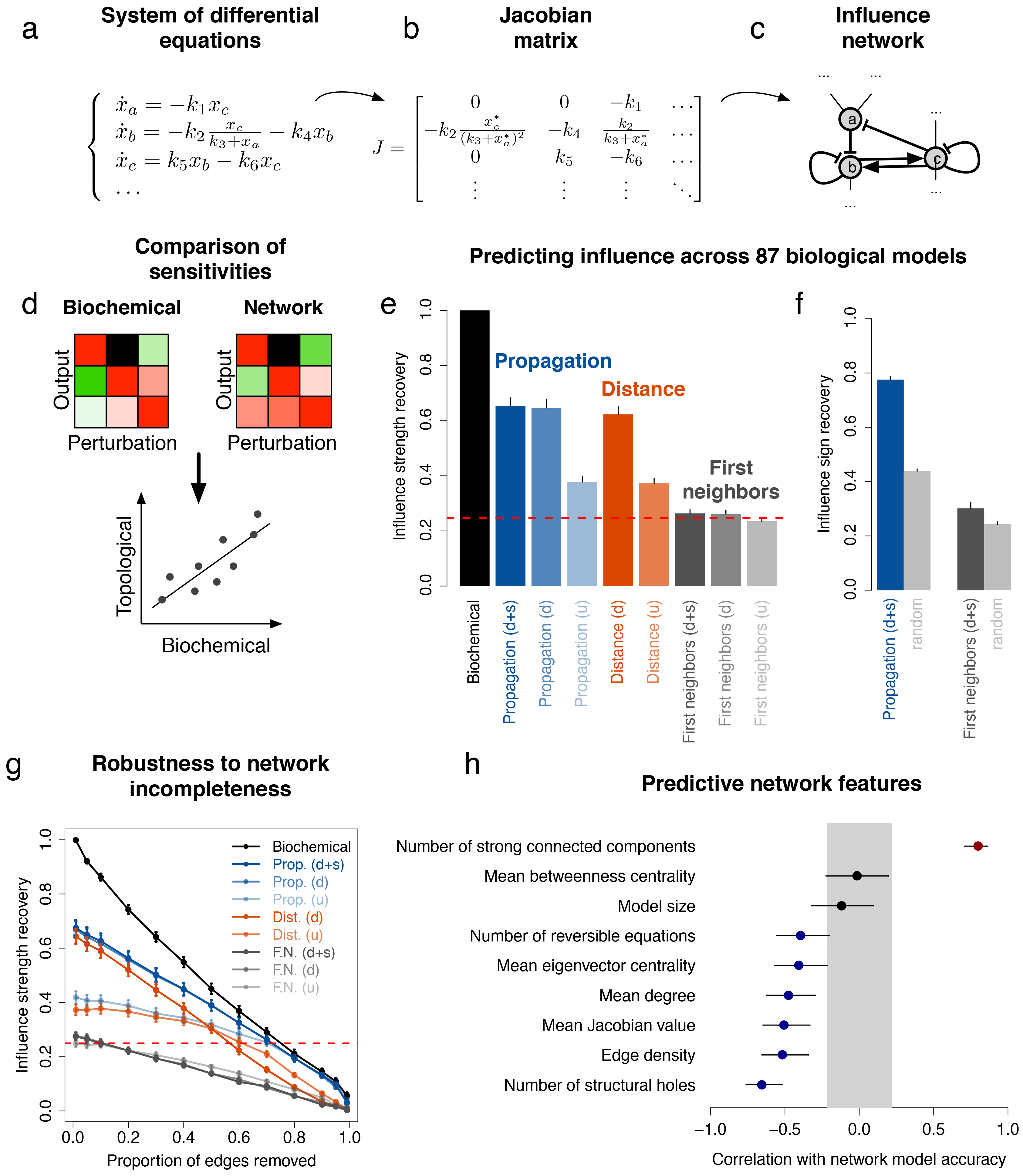
Topology predicts influence patterns in various biological networks. a. We show a set of example differential equations describing biochemical dynamics. These equations involve the different variables from the model capturing the underlying biology of the problem. b. We derive a Jacobian matrix *J* by perturbing the differential equations around their steady-state *x*\*. c. We convert the equations into an “influence network” where a link from species *i* to *j* is created if *i* changes *j* concentration. This corresponds to the sign of the Jacobian matrix (see Methods). d. Schematic representation of the workflow we used to assess the ability of different models to predict the influence patterns. The calculation of the sensitivity matrices is explained in Methods. We use the Spearman correlation coefficient to compare them with the biochemical, “ground truth” sensitivity matrix. e. Bar plot showing the accuracy of different network models in predicting the influence patterns across 87 models from BioModels. We compare sensitivity matrices of different models to the biochemical sensitivity matrix using Spearman correlation, and we average the resulting correlations over all models. Errors bars correspond to standard error. Red dashed line shows two standard deviation of the random expectation averaged over all models. f. Bar plot showing the accuracy of the signed network models in predicting the influence sign across 87 models from BioModels. Errors bars correspond to standard error. Gray bars show random expectation. g. Accuracy of the network models as a function of the proportion of links removed, averaged over the 87 models. Model names are abbreviated as follows: “prop.” for propagation, “dist.” for distance and “F.N.” for First Neighbors. Errors bars show standard error. h. Correlation between properties (names on the left) and the propagation (d+s) model accuracy across 87 BioModels. Gray area shows the random expectation.

We implemented 87 models from BioModels (Table S1), selected with the criterion that the largest connected component contains at least 10 species. For each model, biochemical and DYNAMO sensitivities are computed as follows (see Methods for additional details). For biochemical models, the sensitivity matrix is obtained via (17, 27): 
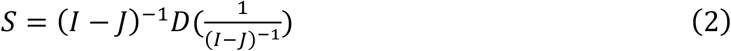
where *I* is the identity matrix and *J* denotes the Jacobian matrix of the system around steady-state (see Methods). For network models, we start from spreading models (i)-(iii) from the DYNAMO model family (Figure 1c-e). To compute the influence of a node on the other nodes in the network, we start by assigning all nodes a weight zero. Given a perturbation in node *i* (weight 1), the influence propagates to other nodes *j* in the network, changing their weights according to various proposed models. The matrix of influence for any pair (*i*, *j*) constitutes the sensitivity matrix. The models are as following: 
i. *Propagation*: We extend the PRINCE methodology to the case of directed and signed networks (29) (see Methods). Noting with *W* the adjacency matrix, we define the diagonal matrix *D*_1_ such that *D*_1_(*i*, *i*) is the sum of the absolute values of row *i* of *W* and the matrix *D*_2_ such that *D*_2_(*i*, *i*) is the sum of the absolute values of column *i* of *W.* We then compute the normalized propagation weights *W* ′=*D*_1_^−1/2^*WD*_2_^−1/2^ and the sensitivity matrix as 
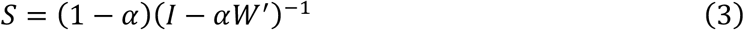
where *α* = 0.9 is a parameter characterizing the propagation strength. This sensitivity matrix corresponds to the spread of a perturbation of weight 1 to the rest of the network.
ii. *Distance:* We assume that influence propagates to all nodes in the same connected component as node *i*. In the directed case, propagation is limited to outgoing links. Weights decrease with distance *d* as 1/(1 + *d*). We note that there is no “signed” case for the distance model. Indeed, there are in theory several shortest paths of the same length joining any two nodes, and it is unclear which one to choose and how to carry the edge signs from the source to the target.
iii. *First neighbors*: We assume that influence propagates only to the direct neighbors of *i,* setting their weights to 1 (or −1 for a negative interaction). For directed networks, only outgoing links are considered.

The sensitivity matrices are compared to the one predicted from the full biochemical model using Spearman correlation (Figure 2d and Methods). This non-parametric measure compares the rank of the sensitivities, not their raw values, thus assessing if the relative strength of perturbations is conserved across models. We use the absolute value of the sensitivities as we focus on recovering the strength of perturbations, not their sign. Figure 2e summarizes the obtained correlations averaged over all 87 biochemical models (see Figure S1 for full distributions), documenting a gradual decrease of the accuracy with the decreasing complexity of the network models. We find that the propagation model, which relies on topological information only, achieves 66% of accuracy (i.e a Spearman correlation of *ρ* = 0.66) in predicting the influence patterns when the network includes direction and sign of the links. This is only slightly better than in the unsigned case, though not significantly (65%, p=0.4 under Student t-test), but a dramatic improvement over the undirected case (40%, p=5.1e-13), indicating that capturing the direction of flow is essential for predictive accuracy. Interestingly, the simpler distance model on a directed network shows comparable accuracy to the best propagation model on a directed signed network (63%). Again, the accuracy greatly decreases in the undirected case to a level similar to the undirected propagation model (36%). Finally, simple first-neighbors models achieve up to ~27% accuracy, a value close to but higher than the random expectation (Figure 2e, dashed red line).

Next we explore whether the signed DYNAMO models can correctly predict the signs of the perturbations. Such signs indicate whether the increase in a species concentration causes another species to be up-or down-regulated. This is important as many measurements report sets of down-or up-regulated genes. We therefore compute the proportion of accurate sign predictions using the signed propagation and first neighbor models (Figure 2f). The results show a similar trend with improvement over the influence strength case, with 78% accuracy (p<1e-16) for the propagation model and 33% (p=1.7e-5) for the first neighbors model.

Overall, we show that topology accounts for 2/3 of the accuracy budget of dynamical models when predicting perturbation patterns. In particular, we find that the simple distance-based model has similar performance as the top performing propagation model.

### 3. Robustness to network incompleteness

While Figure 2f confirms the importance of topology in retrieving biochemical influence patterns, it is unclear to what extent the results hold if the underlying interactome is incomplete. Indeed, high-throughput methods cover less than 20% of all potential pairwise protein interactions in the human cell (7). Despite the gradually increasing coverage (1, 36), we can expect to deal with incomplete models for many years to come. This prompts us to address the robustness of our approach to link removal. Since all approaches inherently rely on the Jacobian matrix of the system, removing a non-zero entry is equivalent to removing a link. We show in Figure 2g the average accuracy of the DYNAMO models in retrieving the original biochemical model sensitivities when removing an increasing proportion of links (i.e an increasing proportion of entries from the original Jacobian matrix). We observe two different behaviors. For directed models the accuracy decreases linearly, while for undirected models it has a concave shape, decreasing slowly initially then more rapidly with additional link removal. This can be understood by realizing that many models have a substantial fraction of reversible equations, modeled as two links of opposite direction between two nodes. In the undirected case, removing one of those two links does not change the network, making these models therefore more robust to link removal. Moreover, we find that at 50% incompleteness, the propagation and biochemical models have similar accuracies, with the biochemical model still slightly better than the propagation one (45% vs 39%). This demonstrates that with the current level of incompleteness of biochemical networks topological models are competitive with more complex “kinetics-aware” models.

### 4. Network features underlying accurate influence prediction

Given the differences in size and scope across the 87 biological models (Table S1), next we investigate what network characteristics contribute to higher prediction accuracy. For this we measured the correlation between various quantities and the directed signed propagation model accuracy across 87 biological models (Figure 2h). The gray area specifies the 95% Confidence Interval. We find that the model size has no effect on accuracy, while the dynamics does: the presence of very high Jacobian values, corresponding to fast reactions, lead to smaller accuracies. This stems from the fact that such outliers in the Jacobian matrix can outweigh the other links and lead to faster propagation across selected links in the biochemical model, a feature that cannot be captured by the network topology alone. We also find that the proportion of reversible equations negatively impacts the accuracy. Indeed, directedness does not offer an advantage for network models for BioModels dominated by reversible equations, and the directed propagation model closes the gap with the less accurate undirected model. This result is supported by the finding that higher accuracies can be reached for networks that can be decomposed into a large number of strongly connected components (SCCs), that is subgraphs for which every node is reachable from every other node. When filtering out BioModels with only one SCC, we observe significant improvement of the DYNAMO accuracies, reaching to ~80% accuracy (Figure S3). We show in Figure S4 two example BioModels with respectively N=1 and N=5 SCCs. The network with 1 SCC is dense and poorly modular, while the one with 5 SCCs is sparser and displays chain-like structures. Supporting such structures, we find higher accuracies for networks that do not display clear hubs (low average degree, eigenvector centrality, and proportion of structural holes), and are sparse (low link density). Finally, since the link weight plays a role through the Jacobian, we would expect that link betweenness centrality should similarly matter, but we find no significant correlation. Overall, we find that topological models reach up to 80% accuracy for biological models with certain network characteristics pertaining to sparsity and modularity.

### 5. Comparison with a Normalized-Hill model

While our DYNAMO framework encompasses a broad range of topological models, it does not include any combinatorial information and we do not model how entities combine when influencing a node - we consider these combinations to be “OR gates”, i.e. additive functions. Here we compare our simplified DYNAMO models to the kinetic-agnostic Boolean-like Normalized Hill Model (NHM, see methods) proposed by (20), that encapsulates such combinatorial features from the original biochemical model (Figure 3a). The NHM represents the dynamics by sigmoidal activation or inhibition functions parametrized through 3 shape parameters, and allows for multiplicative inputs (“AND gates”). Such features, absent from our DYNAMO framework, are designed to offer more realistic insights into the full biochemical model. Our goal is to quantify the residual accuracy in the DYNAMO framework resulting from ignoring combinatorial inputs. The NHM has previously been applied to the beta-adrenergic signaling pathway, a well-studied signaling network regulating cardiac myocyte contractility and involved in cardiac hypertrophy and heart failure (Figure 3b). The full biochemical model (37) contains 87 model parameters, characterizing the interactions between 25 species via 33 links (37). The biochemical model was approximated by the NHM, finding that the resulting sensitivity matrix shows a good correspondence to the full biochemical model (20). Moreover, the NHM was refined to reproduce accurate temporal dynamics of several key proteins by fitting 11 parameters to full time-course data obtained with the full biochemical model (referred hereafter as the “NHM fit” model).

**Figure 3.**
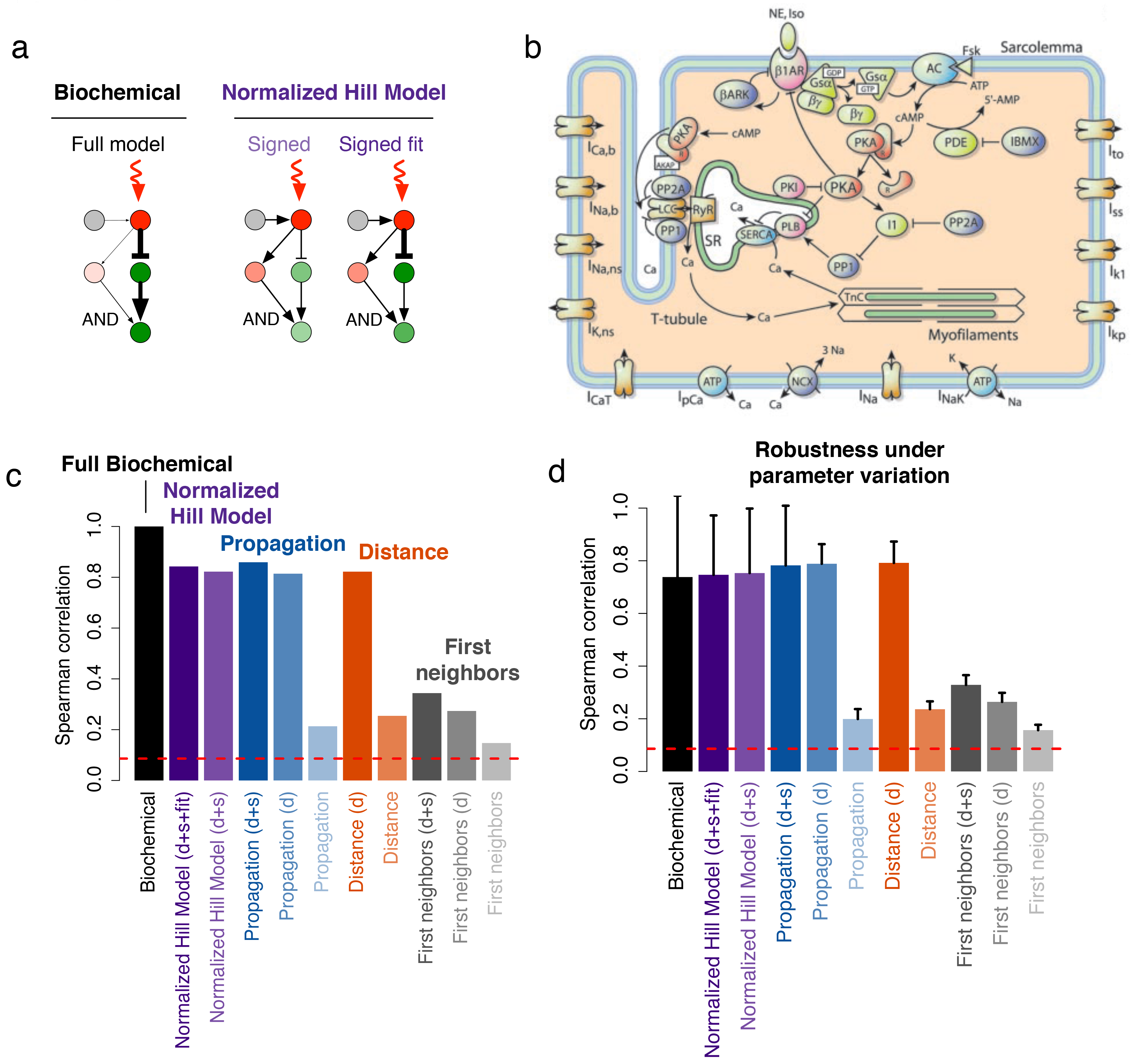
Topology predicts influence in a signaling network. **a.** Schematic representations depicting the propagation of a perturbation in a biological network according to the biochemical and the Normalized Hill Model. “Signed-fit” correspond to the fitting of 11 parameters from this model to reproduce key quantitative time-course behaviors, as described in (20). **b.** Beta-adrenergic signaling network (37). The full biochemical model contains 87 parameters. **c.** Same as Figure 2e. **d.** Robustness of the Spearman correlation measure for the beta adrenergic pathway under parameter perturbation (see Methods). We show the average correlation between any two perturbed biochemical sensitivity matrices generated (black bars), or the average correlation between each network model and the perturbed biochemical sensitivity matrices (other bars). Error bars show standard error. Interestingly, we see no difference between a typical perturbed biochemical model and a network model.

Here we apply our DYNAMO framework to the beta-adrenergic network and compare the resulting sensitivity matrices to the original biochemical one (Figure 3c and sensitivity matrices shown in Figure S5). As previously, we observe a gradual decrease of the accuracy of influence patterns with the decreasing complexity of the network models. Interestingly, the directed propagation and distance models show accuracies of ~80%, similar to the accuracies for both the NHM and NHM fit models.

We then explore the robustness of these results to random variations in the biochemical parameters (see Methods and Figure 3d). For this we generate perturbed biochemical models by multiplying all parameters by a factor randomly chosen between ½ and 2. The resulting sensitivity matrices are then used as ground truth biochemical model to compare with the DYNAMO models. We observe that the accuracy of the network models is mostly unchanged when comparing to the perturbed biochemical models (Figure 3d). However, when comparing the perturbed biochemical model sensitivities with one another, we observe an average accuracy of ~80%, similar to that of the best network models (NHM, propagation and distance models). This indicates that biochemical models with poorly measured kinetic parameters (up to 2-fold variation) are as accurate as topological models. This exhibits the utility of such simple topological models in the context where obtaining precise kinetic information would require important experimental investment.

### 6. Topology predicts physiological and phenotypic perturbations

While perturbation patterns are of general interest for assessing the quality of our models, testing the true value of the network topology-based modeling framework needs experimental validation. To test the accuracy of the DYNAMO models against experimental observations, we focused on the chemotaxis network in bacteria (Figure 4a) for which experimental data is available (38) (see Methods). This model is part of BioModels, and the DYNAMO models capture its dynamics with a 90% accuracy (Figure 4b). The experimental dataset consists of knockouts and overexpression assays of 6 genes of the chemotaxis network and their combinations, followed by observation of the change in expression of other genes from the network. In addition, the experiments also report changes in bias, a phenotypic quantity determined by the ratio of multiple biochemical species concentrations and capturing the exploratory behavior during chemotaxis (see Methods and Figure 4c). In Figure 4d,e we compare the experimental observations (left columns) to the predictions from the propagation model on the signed directed network (right columns) under several assays. We focus on the accuracy in retrieving the correct sign of the observed perturbations. We observe that the network model predicts the observed sign of the perturbations in 86% of the cases for gene expression changes (13 out of 15 cases, p=4.9e-4 under binomial test, Figure 4d). Moreover, it predicts phenotypic changes with 75% accuracy (9 out of 12 cases, p=0.019, Figure 4e), demonstrating the value of topological models in predicting physiologically relevant biological outcomes. Taken together, these results demonstrate that the precision holds when using data from experimental perturbations.

**Figure 4.**
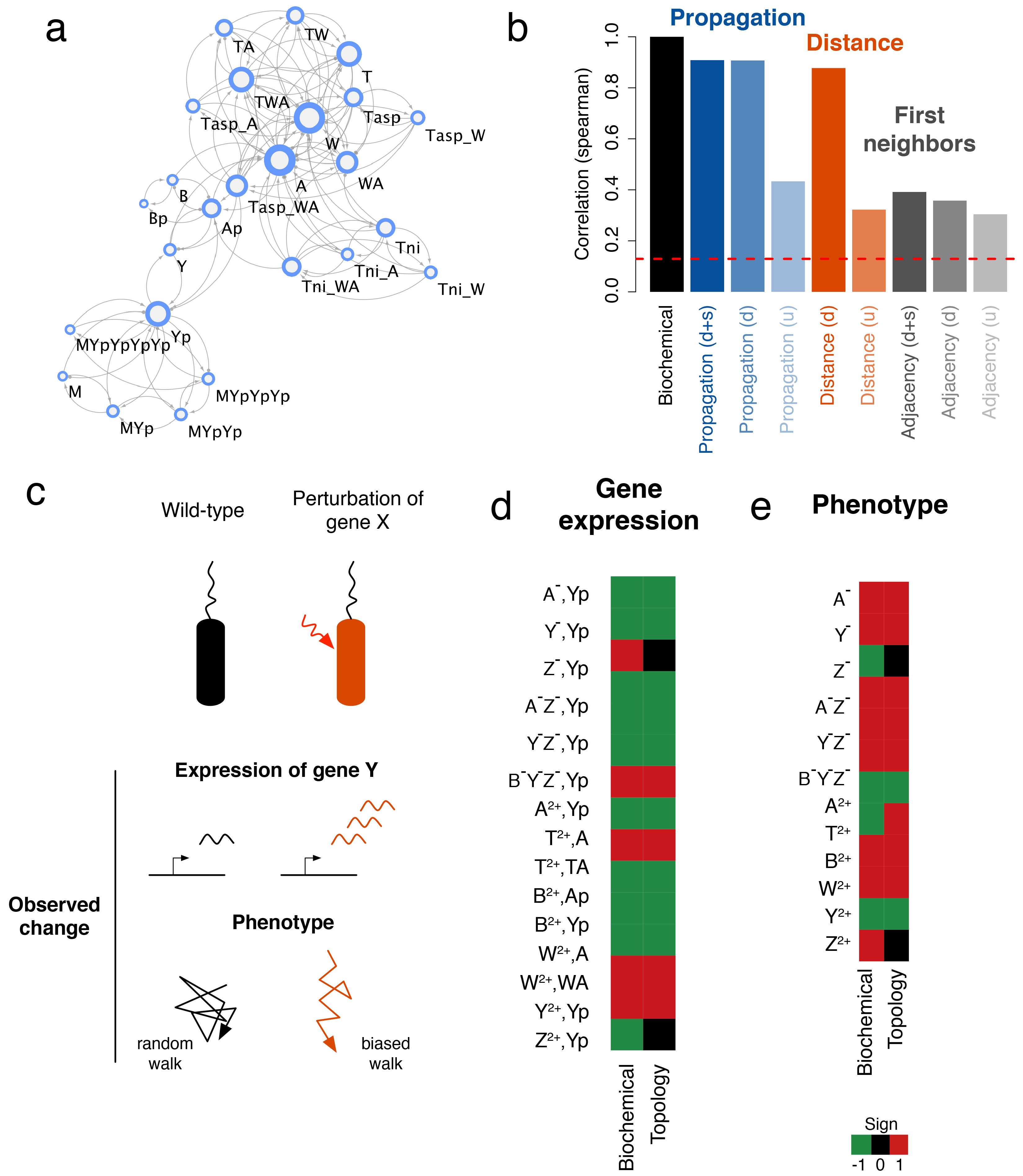
Experimental validation. We compare the results of our influence predictions to experimentally validated perturbations in the chemotactic pathway in bacteria (38). **a.** Chemotaxis network obtained from BioModels. Arrows indicate a positive link weight, and circles a negative one. For each node we indicate the corresponding biochemical species. A detailed description of the model can be found in (38). **b.** Same as Figure 2e for the chemotaxis model. **c.** Schematic representation of the experiments, comparing wild-type bacteria (left, black) with bacteria perturbed with a gene X knock-out or overrepresentation (right, red). The variables of interest consist of gene Y expression change (increased expression corresponding to more mRNA produced, in red), and change in walk bias compare to random walk, in this case an increase (red), corresponding to a more directed exploration behavior. **d.** We compare differential expression from perturbations experiments to predictions from the propagation model (see Methods). The color code indicates the sign of the observed expression change: green, negative, black, no change, red, positive. The names of the rows show the perturbed species (left) and the measured species (right) separated by a comma. Perturbed species consist of single null mutants (*A*^−^, *Y*^−^, *Z*^−^), multiple null mutants (*A*^−^*Z*^−^, *Y*^−^*Z*^−^), and overproduction mutants (*A*^2+^, *T*^2+^, *B*^2+^, *W*^2+^, *Y*^2+^, *Z*^2+^). Measured species consist of *Y*_*p*_, *A*, *TA*, *Ap*, and *WA*. Results agree in 86% of the cases. **e.** We assess phenotypic effect by looking at the sign of the change in bias (see Methods). Row names show perturbed species. Results agree 75% of the cases.

## Discussion

While the coverage of the physical interactions underlying biological networks has increased considerably in the past decade, we continue to lack accurate and comprehensive data on the kinetic parameters determining the dynamics of each individual process. We must therefore evaluate the predictive potential of purely topological models, quantifying their ability to unlock quantitative insights on physiologically relevant processes. In this work, we proposed a systematic DYNAMO framework to measure the loss in predictive power when we lack the kinetic parameters in a biological network. We concentrated on the patterns characterizing the spread of perturbations of selected biochemical species, and its impact on all other species in the network. Such patterns are of direct interest as we seek to understand changes in gene expression patterns induced by disease-causing mutations. We used detailed dynamical biochemical models derived from the literature to estimate the accuracy of the topological models in retrieving the perturbation patterns characterizing the full dynamics. Interestingly, we find that a propagation model on a topology that captures the direction and sign of the interactions can account for ~65% of the full perturbation patterns across all models, an accuracy that can reach 80% for models with certain network characteristics. Furthermore, this model shows only mild improvement over a simpler distance-based model on a directed network. This is important since network distance in the interactome has been used extensively to predict disease similarity between disease genes (7) and drug-disease association between a set of drug targets and disease genes (30). We also find that a simpler first neighbor model or complex models that lack directionality, though offering smaller accuracy, still carry predictive potential compared to a random reference frame. Moreover, the beta-adrenergic example demonstrated that the best DYNAMO topological models can be as informative as slightly perturbed biochemical models where kinetic parameters would carry measurement noise. They also offer an accuracy comparable to models with more complex non-linear dynamics, key parameters fit and multiplicative input functions.

A usual concern raised about the predictive power of biochemical networks is that the interactome is far from complete. We therefore addressed the robustness of our approach to link removal, finding that the DYNAMO models are in general more robust than the biochemical model to link removal, and that the propagation model has comparable accuracy to the biochemical model at 50% incompleteness. Therefore, we find that the kinetic-agnostic DYNAMO models are as predictive as the full biochemical models in the context of incomplete interactome.

We also explored which network characteristics offer greater accuracy for the DYNAMO framework. This is important as it allows to know before any kinetic parameters have been measured whether that information would lead to a drastic improvement over a topological model. The analysis indicates that networks that can be decoupled into many strong connected components (i.e many chain-like structures) and are in general sparse (low degree nodes and link density) lead to higher DYNAMO accuracies.

Finally, exploring the topological models’ ability to predict observed outcomes from experimental perturbations in the chemotaxis pathway, we find a 75% to 86% accuracy, a result in agreement with the previous accuracies computed for the full perturbation patterns.

This work has important implications for our understanding of the role of the different modeling frameworks currently in use in network biology and medicine. The ability to extract information on the spread of perturbations from an accurate knowledge of the topology of biological networks will be of great value for drug development. In particular, the linearity of the equation used to predict perturbation patterns in the propagation model makes it straightforward to explore any combination of perturbations, therefore paving the way for a better understanding of drug combinations and improved therapies. Overall, our findings indicate that the lack of large scale measurements of the kinetic parameters may not prevent network biology to offer quantitative and accurate predictions on perturbation processes. On the contrary, focusing on topology appears well justified as it holds the largest share of the descriptive power in this endeavor.

## Methods

### BioModels database

Models from the BioModels database were downloaded in bulk from the BioModels ftp website (ftp://ftp.ebi.ac.uk/pub/databases/biomodels/). We used the BioModels_Database-r30_pub-sbml_files dataset (ftp://ftp.ebi.ac.uk/pub/databases/biomodels/releases/2016-05-10/). SBML files were processed using libsbml matlab library to extract the dynamical equations (34).

### Derivation of the influence network

We converted the reactions from BioModels SBML files to reaction networks using libSBML (34). We first extracted the differential equations describing the dynamical models in the form 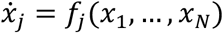 where *N* describes the number of species in the model and *x*_*i*_ is the concentration of species *i*. These equations are in turn converted to an “influence network” where a link from species *i* to j is created if *i* is in the differential equation describing the evolution of *j*, i.e if 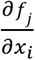 is not identically zero. The sign of the link is the sign of 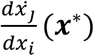 where *x*\* is the steady state vector of species concentration. In other words, the influence network is *sign*(*J*^*T*^(*x*\*)) where *J*^*T*^ is the transpose of the Jacobian describing the system. Models were integrated using the ode23tb function from Matlab R2016a and the Jacobian was computed using finite difference method with step size 1e-3.

### Computation of biochemical sensitivities

The biochemical sensitivities are computed following (17, 27). Writing the dynamical equations as 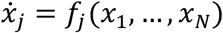 where *N* is the number of species in the model and *x*_*i*_ is the concentration of species *i*, we define the Jacobian by 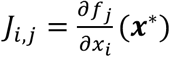 where *x*\* is the steady-state of the system (Figure 2b). The Jacobian captures the impact that a small perturbation in *x*_*i*_ has on the value of *x*_*j*_, providing a quantitative measure for the influence of *i* on the activity of *j*. The partial derivative implies that no other node activity has changed, so that the Jacobian only captures direct interactions. To account for indirect interactions we further define the *sensitivity matrix* by 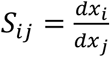 Here, the full derivative implies that all nodes are allowed to change in response to *j*’S perturbation, hence indirect effects are also accounted for. The two matrices are linked by the following equations (17, 27): 
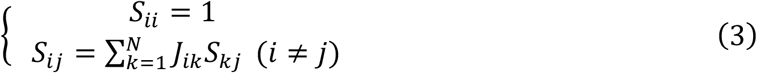

These equations can be simplified by noting that *S* = *JS* off the diagonal, so that we can find a diagonal matrix Δ such that *S* = *JS* + Δ. This leads to *S* = (*I* − *J*)^−1^Δ where *I* is the identity matrix. Finally, using *S*_*ii*_ = 1, we find that 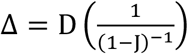, where *D*(.) is the diagonal operator and the operator / denotes element-wise division. Taken together, this leads to the equation (2) in the main text.

### Network models of influence propagation

We use models (i)-(iii) described in the text to predict influence propagation in a network (see Figure 1c-e). For the propagation model, we extended the PRINCE methodology to the case of a directed network by using in-and out-degree in the normalization process (see main text). For the signed network, we used the same normalization as for the signed directed adjacency matrix, ensuring the convergence of the unsigned random walk. To better understand the extension to the signed case, we return to the equation of the underlying perturbation (29): 
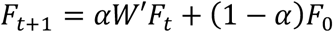
where *F*_*t*_ represents the vector of perturbation at time t, (1 − *α*) is the return probability, *W* ′ is the normalized signed directed adjacency matrix, and *F*_0_ is the initial perturbation. A perturbation spreads to neighboring nodes with a probability given by the absolute value of the corresponding element in *W* ′ and is multiplied by the sign of this element. Accordingly, sensitivities can be thought of as a weighted sum over all network paths between a perturbed node and an impacted node, with each path carrying a positive (negative) weight if the path contains an even (odd) number of negative signs. As such, the absolute values of the sensitivities of the directed signed diffusion model are smaller or equal to the sensitivities of the directed diffusion model, ensuring convergence.

Note that in the propagation mode, the variation of *α* changes the diffusion capability. However, we observed no significant change in the influence strength recovery across 87 BioModels when varying *α* (Figure S2, panel A). Indeed, we are interested in the ranks of the observed sensitivities (the most vs the least perturbed species), which is why we use the non-parametric Spearman rank correlation. We show in the case of the chemotaxis pathway that while the change of *α* alters the values predicted by the propagation model (panels B, C), it has no effect on the rank of the model sensitivities (panel D, E), leaving the Spearman correlation almost unchanged (panel A).

In the case of the beta-adrenergic pathway, we use the Normalized Hill Function system developed in (20). This model includes a simple non-linear dynamics and multiplicative inputs on select cases as informed from the literature. Species interactions were defined with normalized activating or inhibiting Hill functions (*f*_*act*_ or *f*_*inhib*_) and pathway crosstalk was implemented using logical AND and OR operations: “*f*(*x*)*f*(*y*)” and “*f*(*x*) + *f*(*y*) − *f*(*x*)*f*(*y*)”, respectively. The normalized activating or inhibiting Hill functions have the form: 
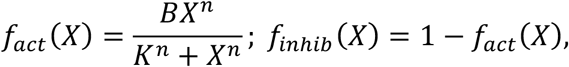
where *B* and *K* are constrained such that *f*_*act*_(0) = 0, *f*_*act*_(*EC*50) = 0.5, and *f*_*act*_(1) = 1. From these constraints, we derive: 
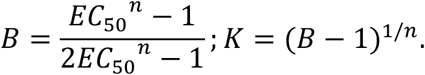

As default parameters we used *EC*_50_ = 0.5, *n* = 1.4, and *τ* = 1.

For the NHM fit model, we used the refined NHM model from (20) where several parameters in the normalized-Hill model are fitted to time-course data from the biochemical model using a nonlinear least squares optimization algorithm. Eleven parameters were adjusted to fit model predictions, allowing for more similar signaling dynamics and comparable peak fractional activities of species *GS*_*α*_ and PLB compared with the biochemical model (20).

### Comparison of sensitivity matrices

To evaluate the accuracy of each model, we computed the Spearman correlation between their sensitivity matrix and the one obtained with the full biochemical model. For unsigned networks, we use the absolute value of the sensitivity matrix.

### Robustness to network incompleteness

To model network incompleteness, for each model we successively removed entries of the Jacobian (the links of the influence network). A proportion of links removed was computed as the number of entries removed divided by the total number of non-zero initial entries. The links were chosen at random and the resulting incomplete Jacobian was finally used to compute the new sensitivity matrix as well as the topological models for DYNAMO. The random process was iterated 20 times.

### Comparison with experimental data

For the chemotaxis model, experimental data was obtained from (38). We built the influence network from the corresponding BioModel BIOMD0000000404. The propagation model was used on the directed, signed network. We first explored whether the propagation model retrieved the direction of change of the expression of gene *i* caused by a mutation or overexpression of gene *j* for the experimentally investigated cases. To do so, we retrieved the signs of elements (*i*, *j*) from the predicted sensitivity matrix and compared them to the experimental ones. In the case of mutations, the negative of the sensitivity matrix was used. In the case of multiple mutations *j*_1_ …*j*_*N*_, the corresponding columns of the sensitivity matrix were summed and the resulting vector was used to retrieve the sign of the perturbation of interest. We then tested whether our model could retrieve the change of a more complex phenotypic quantity, namely the bias, determined by the ratio of the following biochemical species concentrations: 
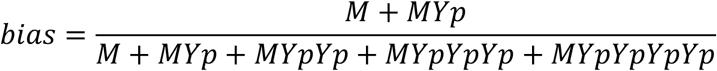

Following the experiments of Ref. (38), we explored the impact of perturbations on the bias by computing the sign of its change after perturbation. Noting 
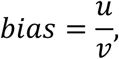
we derived that for a perturbation *dx*_*j*_ 
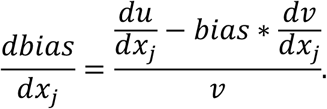

Therefore the sign of the change of bias is given by 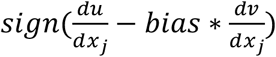, where *bias* = 0.7 is given by the full biochemical model (38), and 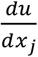 and 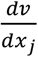 are obtained from sensitivity matrix of the propagation model.

### Robustness of Sensitivity matrices

We computed the robustness of the Spearman correlation measure under parameter variation (initial conditions and kinetic parameters). Parameters from the biochemical model were multiplied by a factor randomly chosen between 1/2 and 2, and the corresponding sensitivity matrix was computed. We repeated the process to obtain 100 sensitivity matrices, discarding cases where the dynamical models diverged, representing 35% of the trials.

### Code availability

The algorithms used in that work are made available on https://github.com/msantolini/dynamo.

## Acknowledgements

We thank B. Barzel, L. Blondel, I. Kovacs and A. Krishna for careful reading and helpful comments about the manuscript. This work was supported by the National Institutes of Health through National Heart, Lung, and Blood Institute Grant 1P01HL132825-01 and Centers of Excellence in Genomic Science Grant P50HG004233.

## References

1. Rolland T, et al. (2014) A proteome-scale map of the human interactome network. Cell 159(5):1212–1226.

2. Venkatesan K, et al. (2009) An empirical framework for binary interactome mapping. Nat Methods 6(1):83–90.

3. Buchanan M, Caldarelli G, De Los Rios P, Rao F, & Vendruscolo M (2010) Networks in cell biology (Cambridge University Press, Cambridge; New York) pp x, 271 p.

4. Barabasi AL & Oltvai ZN (2004) Network biology: understanding the cell’s functional organization. Nat Rev Genet 5(2):101–113.

5. Barabasi AL, Gulbahce N, & Loscalzo J (2011) Network medicine: a network-based approach to human disease. Nat Rev Genet 12(1):56–68.

6. Vidal M, Cusick ME, & Barabasi AL (2011) Interactome networks and human disease. Cell 144(6):986–998.

7. Menche J, et al. (2015) Disease networks. Uncovering disease-disease relationships through the incomplete interactome. Science 347(6224):1257601.

8. Ideker T & Sharan R (2008) Protein networks in disease. Genome Res 18(4):644–652.

9. Clough E & Barrett T (2016) The Gene Expression Omnibus Database. Methods Mol Biol 1418:93–110.

10. Dubitzky W, Southgate J, & Fuss H (2011) Understanding the dynamics of biological systems lessons learned from integrative systems biology. (Springer, New York).

11. Maerkl SJ & Quake SR (2007) A systems approach to measuring the binding energy landscapes of transcription factors. Science 315(5809):233–237.

12. Teusink B, et al. (2000) Can yeast glycolysis be understood in terms of in vitro kinetics of the constituent enzymes? Testing biochemistry. Eur J Biochem 267(17):5313–5329.

13. Mendes P & Kell D (1998) Non-linear optimization of biochemical pathways: applications to metabolic engineering and parameter estimation. Bioinformatics 14(10):869–883.

14. Jaqaman K & Danuser G (2006) Linking data to models: data regression. Nat Rev Mol Cell Biol 7(11):813–819.

15. Rodriguez-Fernandez M, Mendes P, & Banga JR (2006) A hybrid approach for efficient and robust parameter estimation in biochemical pathways. Biosystems 83(2-3):248–265.

16. Brodersen R, Nielsen F, Christiansen JC, & Andersen K (1987) Characterization of binding equilibrium data by a variety of fitted isotherms. Eur J Biochem 169(3):487–495.

17. Barzel B & Barabasi A-L (2013) Universality in network dynamics. in Nature Physics, pp 673–681.

18. Barzel B, Liu YY, & Barabasi AL (2015) Constructing minimal models for complex system dynamics. Nat Commun 6:7186.

19. Davidich MI & Bornholdt S (2013) Boolean network model predicts knockout mutant phenotypes of fission yeast. PLoS One 8(9):e71786.

20. Kraeutler MJ, Soltis AR, & Saucerman JJ (2010) Modeling cardiac beta-adrenergic signaling with normalized-Hill differential equations: comparison with a biochemical model. BMC Syst Biol 4:157.

21. van Gend C & Snoep JL (2008) Systems biology model databases and resources. Essays Biochem 45:223–236.

22. Chelliah V, et al. (2015) BioModels: ten-year anniversary. in Nucleic Acids Res, pp D542–D548.

23. Soltis AR & Saucerman JJ (2011) Robustness portraits of diverse biological networks conserved despite order-of-magnitude parameter uncertainty. in Bioinformatics, pp 2888–2894.

24. Alon U, Surette MG, Barkai N, & Leibler S (1999) Robustness in bacterial chemotaxis. Nature 397(6715):168–171.

25. Gutenkunst RN, et al. (2007) Universally Sloppy Parameter Sensitivities in Systems Biology Models. in PLoS Comput Biol, p e189.

26. Huang B, et al. (2017) Interrogating the topological robustness of gene regulatory circuits by randomization. PLoS Comput Biol 13(3):e1005456.

27. Barzel B & Biham O (2009) Quantifying the connectivity of a network: the network correlation function method. Phys Rev E Stat Nonlin Soft Matter Phys 80(4 Pt 2):046104.

28. Ryall KA, et al. (2012) Network Reconstruction and Systems Analysis of Cardiac Myocyte Hypertrophy Signaling. in Journal of Biological Chemistry, pp 42259–42268.

29. Vanunu O, Magger O, Ruppin E, Shlomi T, & Sharan R (2010) Associating genes and protein complexes with disease via network propagation. PLoS Comput Biol 6(1):e1000641.

30. Guney E, Menche J, Vidal M, & Barabasi AL (2016) Network-based in silico drug efficacy screening. Nat Commun 7:10331.

31. Gulbahce N, et al. (2012) Viral perturbations of host networks reflect disease etiology. PLoS Comput Biol 8(6):e1002531.

32. Oti M, Snel B, Huynen MA, & Brunner HG (2006) Predicting disease genes using protein-protein interactions. J Med Genet 43(8):691–698.

33. Wuchty S (2014) Controllability in protein interaction networks. Proc Natl Acad Sci U S A 111(19):7156–7160.

34. Bornstein BJ, Keating SM, Jouraku A, & Hucka M (2008) LibSBML: an API library for SBML. Bioinformatics 24(6):880–881.

35. Liu Y-Y, Slotine J-J, & Barabasi A-L (2013) Observability of complex systems. in Proc Natl Acad Sci USA, pp 2460–2465.

36. Mosca R, Pons T, Ceol A, Valencia A, & Aloy P (2013) Towards a detailed atlas of protein-protein interactions. Curr Opin Struct Biol 23(6):929–940.

37. Saucerman JJ, Brunton LL, Michailova AP, & McCulloch AD (2003) Modeling beta-adrenergic control of cardiac myocyte contractility in silico. J Biol Chem 278(48):47997–48003.

38. Bray D, Bourret RB, & Simon MI (1993) Computer simulation of the phosphorylation cascade controlling bacterial chemotaxis. Mol Biol Cell 4(5):469–482.

